# The RU486-dependent activation of the GeneSwitch system in adult muscles leads to severe adverse effects in Drosophila

**DOI:** 10.1101/2023.09.07.556712

**Authors:** Maria Paula Zappia, Deena Damschroder, Anton Westacott, Robert J. Wessells, Maxim V. Frolov

## Abstract

Robust genetic systems to control the expression of transgenes in a spatial and temporal manner are a valuable asset for researchers. The GeneSwitch system induced by the drug RU486 has gained widespread use in the Drosophila community. However, some concerns were raised as negative effects were seen depending on the stock, transgene, stage and tissue under study. Here, we characterized the adverse effects triggered by activating the GeneSwitch system in adult muscles using the *MHC-GS-GAL4* driver. When a control, mock *UAS-RNAi* transgene was induced by feeding adult flies with RU486, we found that the overall muscle structure, including myofibrils and mitochondrial shape, was significantly disrupted and led to a significant reduction in the lifespan. Remarkably, lifespan was even shorter when two copies of the driver were used even without the mock *UAS-RNAi* transgene. Thus, researchers should be cautious when interpreting the results given the adverse effects we found when inducing RU486-dependent *MHC-GS-GAL4* in adult muscles. To counter the impact of these effects we recommend setting up additional control groups, such as a mock UAS-RNAi transgene, to validate the findings when using this inducible genetic system, as comparing the phenotypes between RU486-treated and untreated animals could be insufficient.

## Introduction

A wide range of genetic tools have been developed across model organisms to manipulate the expression of genes of interest in a timely and spatial manner. In *Drosophila melanogaster*, the GAL4/UAS system has been broadly used to spatially restrict the expression of the transgene (Brand and Perrimon 1993). Thousands of the UAS transgenes have been generated over the last three decades to modulate the expression of genes of interest and expand the applications of the GAL4/UAS system. Moreover, this system has been modified to temporally control the activation of GAL4 to make it inducible at specific time points during the fly life cycle. One of the inducible GAL4/UAS systems is the “GeneSwitch” system, which consists of a modified progesterone-receptor GAL4 fusion protein that can be activated by the ligand RU486 (mifepristone) (Osterwalder *et al*. 2001; Roman *et al*. 2001). A variety of tissue-specific GAL4 drivers have been generated to be used with the conditional RU486-dependent GAL4/UAS system. For instance, the *MHC-GeneSwitch (GS)-GAL4* is specifically expressed in muscles in the presence of RU486 (Osterwalder *et al*. 2001). The advantage of this system is that the activation of the UAS-transgenes can be restricted to adult stages by feeding adult flies food containing RU486, thus allowing to overcome unwanted effects that may occur earlier in development. We recently successfully used this system to overexpress Sestrin in muscles using *MHC-GS*-*GAL4* and study Sestrin function as a mediator of exercise benefits (Kim *et al*. 2020). However, another study reported alteration in gene expression induced by RU486 in animals carrying *MHC-GS*-*GAL4* (Robles-Murguia *et al*. 2019). Additionally, other works listed the challenges of working with the GeneSwitch system (Poirier *et al*. 2008; Scialo *et al*. 2016). During our experiments, we set up a control group using the mock RNAi, *UAS-mCherry-RNAi*. Surprisingly, we found many defects associated with the control group *MHC-GS>mCherry-RNAi*, thus suggesting that the activation of the GeneSwitch *per se* specifically in the adult muscles could lead to defects in muscle physiology irrespectively of the RNAi transgene. Given the importance of having a reliable induction system and to avoid erroneous conclusions, in this study we aimed to comprehensively characterize the defects induced by the activation of the *MHC-GS-GAL4* driver in adult skeletal muscles in flies.

## Results

### The RU486-dependent activation of the *MHC-GS-GAL4* driver alters the overall muscle structure in adult flies

To thoroughly control the secondary response induced by activating the *MHC-GS* driver with RU486, a control group using the mock RNAi, *UAS-mCherry-RNAi*, was set up. The *MHC-GS>mCherry-RNAi* animals were reared on a regular food diet. Then, in order to induce the GeneSwitch system, 3-day-old adult flies were collected and switched to food containing the RU486 drug (RU + /On). As a control, the vehicle, ethanol, was administered to age-matched flies to keep the system silent (RU - /Off). Animals were aged under these two conditions, RU + and RU -, until 5-, 15- and 25-day-old (Figure 1A). At each timepoint we examined the overall structure of the indirect flight muscles by staining sagittal sections with phalloidin to visualize myofibrils, kettin antibody to label Z-discs on sarcomeres, and ATP5A antibody to reveal mitochondria structure. Interestingly, the *MHC-GS>mCherry-RNAi* RU+ flies showed defects that progressed in their severity over time (Figure 1 B-D). In muscles of 5-day-old control animals fed on RU-food, mitochondria were elongated (Off, Figure 1B). In contrast, about 25% of the 5-day-old flies RU+ (On) contained mitochondria that were rounder (Figure 1B). The phenotype was more prominent in the 15-day-old flies, as roughly 50% of the *MHC-GS>mCherry-RNAi* RU+ animals showed some kind of defect in mitochondria structure. Additionally, there were defects in the organization of myofibrils as revealed by staining using Kettin anitbody (Figure 1C). Finally, 25-day-old flies showed completely disrupted mitochondria and sarcomere units (Figure 1D), which is consistent with a progressive phenotype inducing severe damage in the overall muscle structure.

**Figure 1:**
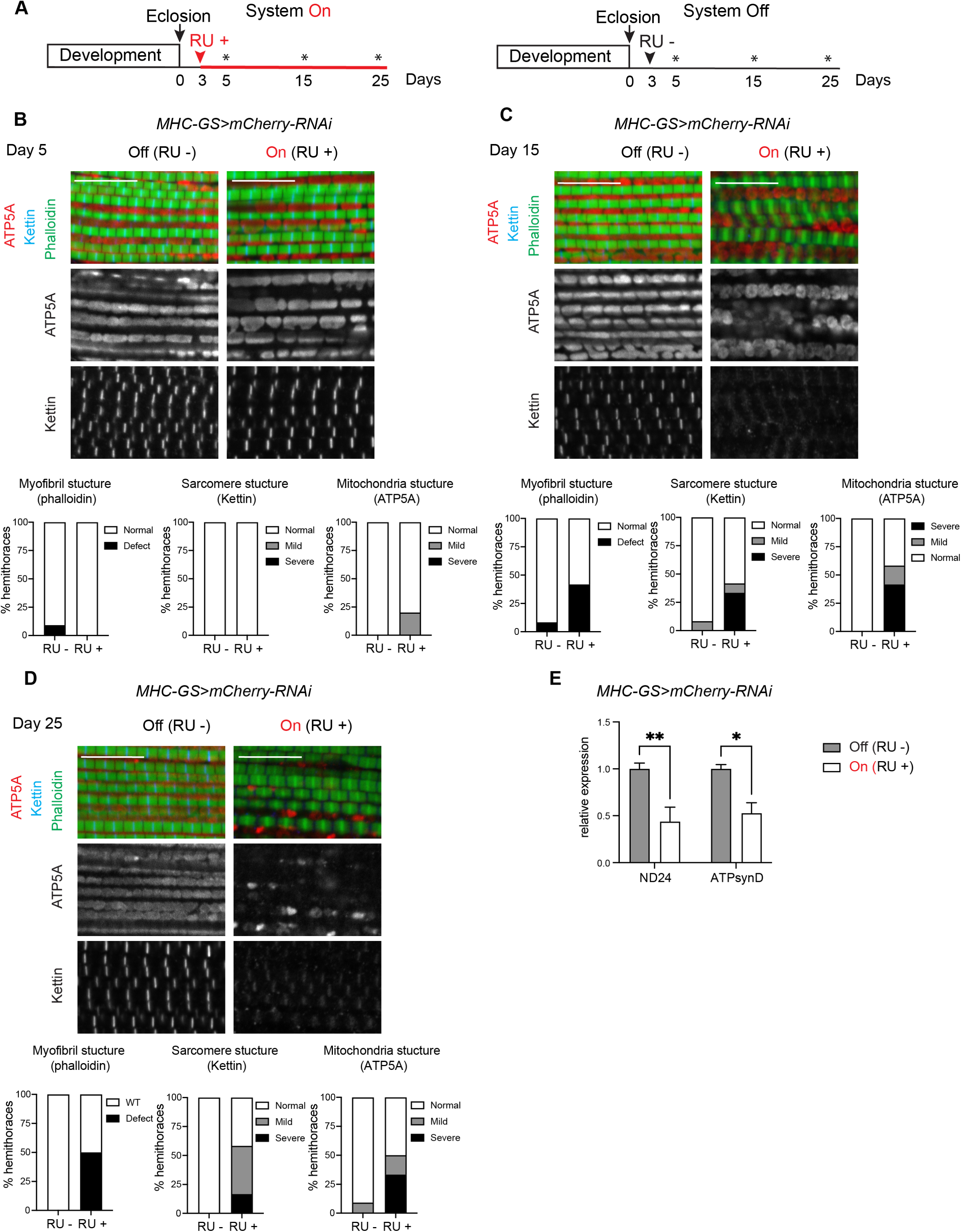
The overall muscle structure in adult flies is altered by the RU486-dependent activation of the *MHC-GS-GAL4* driver A) Timeline of experimental set up. Adult male flies were collected after eclosion (arrow) and 3 days later were treated with RU drug to turn the System On (red arrowhead, RU+, left panel) and vehicle to keep the system Off (black arrowhead, RU-, right panel). Flies were harvested at 5-, 15- and 25-day-old (asterisk). (B-C,D) Confocal section images of flight muscles in a sagittal view. Hemithorax sections of (B) 5-d-old (C) 15-d-old and (D) 25-d-old males stained with Phalloidin (green), anti-kettin (blue), and anti-ATP5A (red). Scale 10 µm. Bottom panels: Quantification of percentage of hemithoraces displaying either normal or defective morphology (mild and severe) for (left) overall myofribril structure, (middle) sarcomere structure, and (right) mitochondria shape (i.e., round). Stacked bars, (B) N = 11-10, (C) N=12, (D) N=11-12 hemithoraces/group. (E) Expression of the nuclear-encoded mitochondrial genes NADH dehydrogenase (ubiquinone) 24 kDa subunit (ND24) and ATP synthase, subunit D (ATPsynD) measured by RT-qPCR in dissected thoraces of 25-day-old adult males. Fold change relative to control. Bar plots with bars, mean ± SD, N = 3 samples/group. Two-way ANOVA, p < 0.01, Bonferroni’s multiple comparisons test. Genotype: *w*/Y*; +; *MHC-GS-GAL4*/*UAS-mCherry-RNAi*

To further characterize these defects, RNA was extracted from thoraces of 25-day-old males and the expression of nuclear encoded-mitochondrial genes were measured by RT-qPCR. The levels of expression of NADH dehydrogenase (ubiquinone) 24 kDa subunit (ND24) and ATP synthase, subunit D (ATPsynD), were significantly reduced when the system was On (Figure 1E). The low expression of the components of mitochondrial complexes I and V are consistent with abnormal mitochondrial morphology, thus suggesting that mitochondria are dysfunctional.

### Activating two copies of *MHC-GS-GAL4* driver results in severe defects

Next, we asked whether the damage in the muscles that was induced by turning On the GeneSwitch system in *MHC-GS>mCherry-RNAi* was caused by either the drug *per se* or one of the genetic elements used, such as *MHC-GS-GAL4* driver and *UAS-mCherry-RNAi* transgene. Therefore, we performed the same experiment as previously mentioned, but this time wild type animals, *w^1118^*, which do not carry any transgene, were used as control. As shown in Figure 2A-C and quantified on the bottom panels, the muscles of *w^1118^* flies RU+ were not altered, as their structure was similar to RU-diet at any given time point.

**Figure 2:**
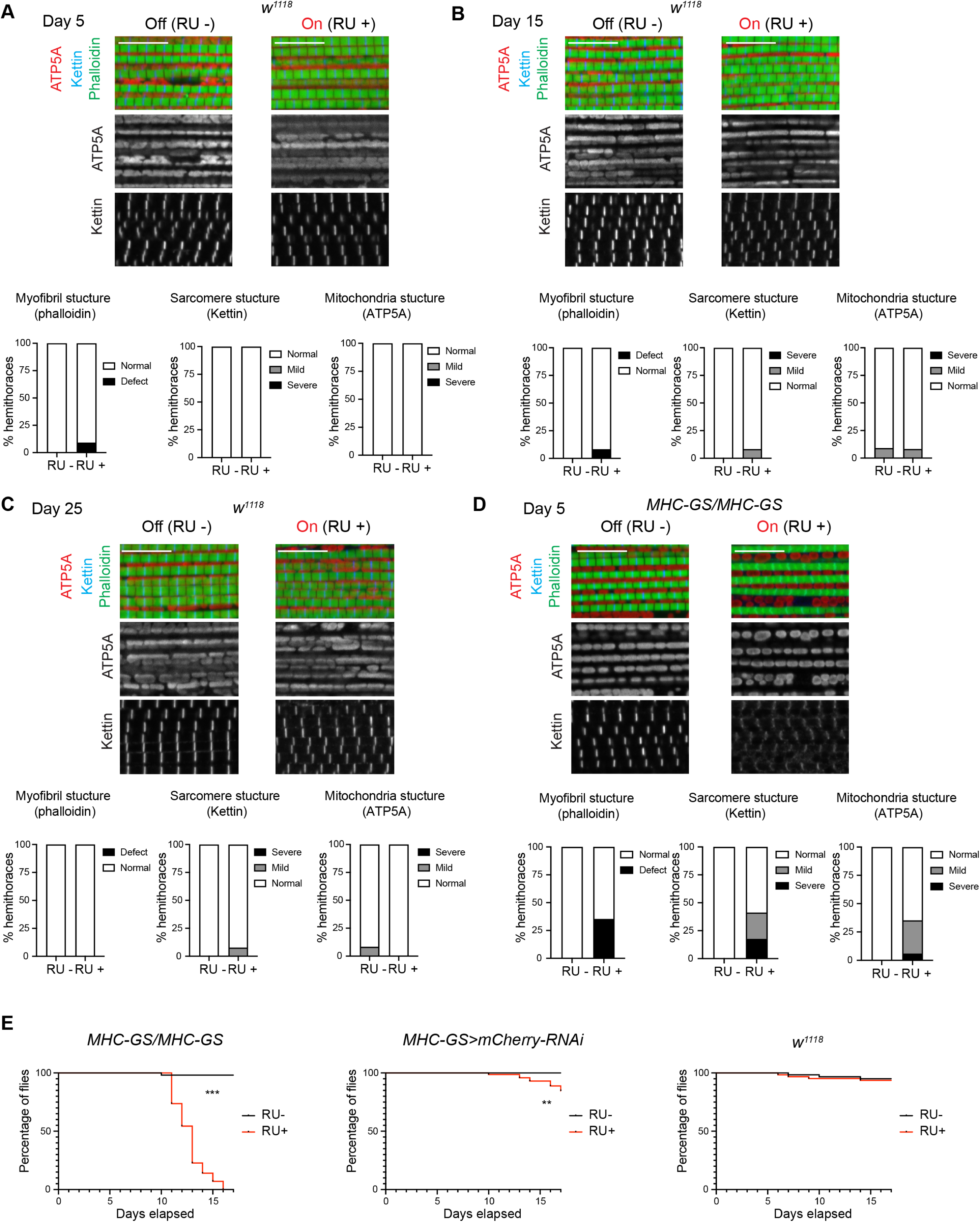
RU486-dependent activation of *MHC-GS-GAL4* homozygote flies is lethal (A-D) Confocal section images of flight muscles in a sagittal view. Hemithorax sections of (A,D) 5-d-old (B) 15-d-old and (C-D) 25-d-old males stained with Phalloidin (green), anti-kettin (blue), and anti-ATP5A (red). Scale 10 µm. Bottom panels: Quantification of percentage of hemithoraces displaying either normal or defective morphology (mild and severe) for (left) overall myofribril structure, (middle) sarcomere structure, and (right) mitochondria shape (i.e., round). Stacked bars, (A) N = 12-11, (B) N=11-12, (C) N=12-13, (D) N=12-17 hemithoraces/group. (E) Kaplan-Meier survival curve. Percentage of viable flies scored scored daily until 17-day-old, Log-rank (Mantel-Cox), *** p<0.001, ** p<0.001, N=53-57 (left panel), N=66-72 (middle panel) and N= 61-63 (right panel). Genotypes: (A-C,D) *w^1118^/Y, (D, E) w*/Y*; +; *MHC-GS-GAL4*/ *MHC-GS-GAL4 (E) w*/Y*; +; *MHC-GS-GAL4*/*UAS-mCherry-RNAi*

Given that the RU+ treatment *per se* was not causing the effects shown in muscles, we hypothesized that either the transgene *MHC-GS-GAL4* or *UAS-mCherryRNAi* was inducing this phenotype. Using a similar experimental approach, we found that at just 5 days old roughly 30% of *MHC-GS-GAL4* homozygous flies, carrying two copies of the driver and no UAS-transgene, showed disrupted myofibrils and round-shape mitochondria (Figure 2D). Interestingly most *MHC-GS-GAL4* homozygous flies died by 15-day old, whereas *w^1118^* did not show any defect when fed with the RU486 drug. Consistently, *MHC-GS>mCherry-RNAi*, carrying one copy of the driver, started to show some decline in viability at 15-day old (Figure 2E).

In sum, our data indicate that the drug itself does not have an effect on flies. However, it seems that the activation of *MHC-GS-GAL4* leads to an alteration in mitochondria and overall muscle structure. This conclusion is further supported by the fact that two copies of this driver resulted in animal lethality as early as 10 days old, indicative of a dose-dependent effect.

### Protein poly-ubiquitination is increased upon activation of the GeneSwitch system in muscles.

The ubiquitin-proteasome system is a major intracellular protein degradation system that plays a critical role in muscle maintenance (Haas *et al*. 2007). Therefore, we tested whether the muscle defects observed in GeneSwitch animals are due to an increased response of the ubiquitin-proteasome system. We examined protein ubiquitination in the adult skeletal muscles of *MHC-GS>mCherry-RNAi* and *w^1118^* by staining thoraces with anti-Ubiquitin antibodies. Poly-ubiquitin aggregates in *MHC-GS>mCherry-RNAi* treated with RU+ were higher than RU-. In contrast, poly-ubiquitination in *w^1118^*did not change in RU+ compared to RU-(Figure 3A-B).

**Figure 3:**
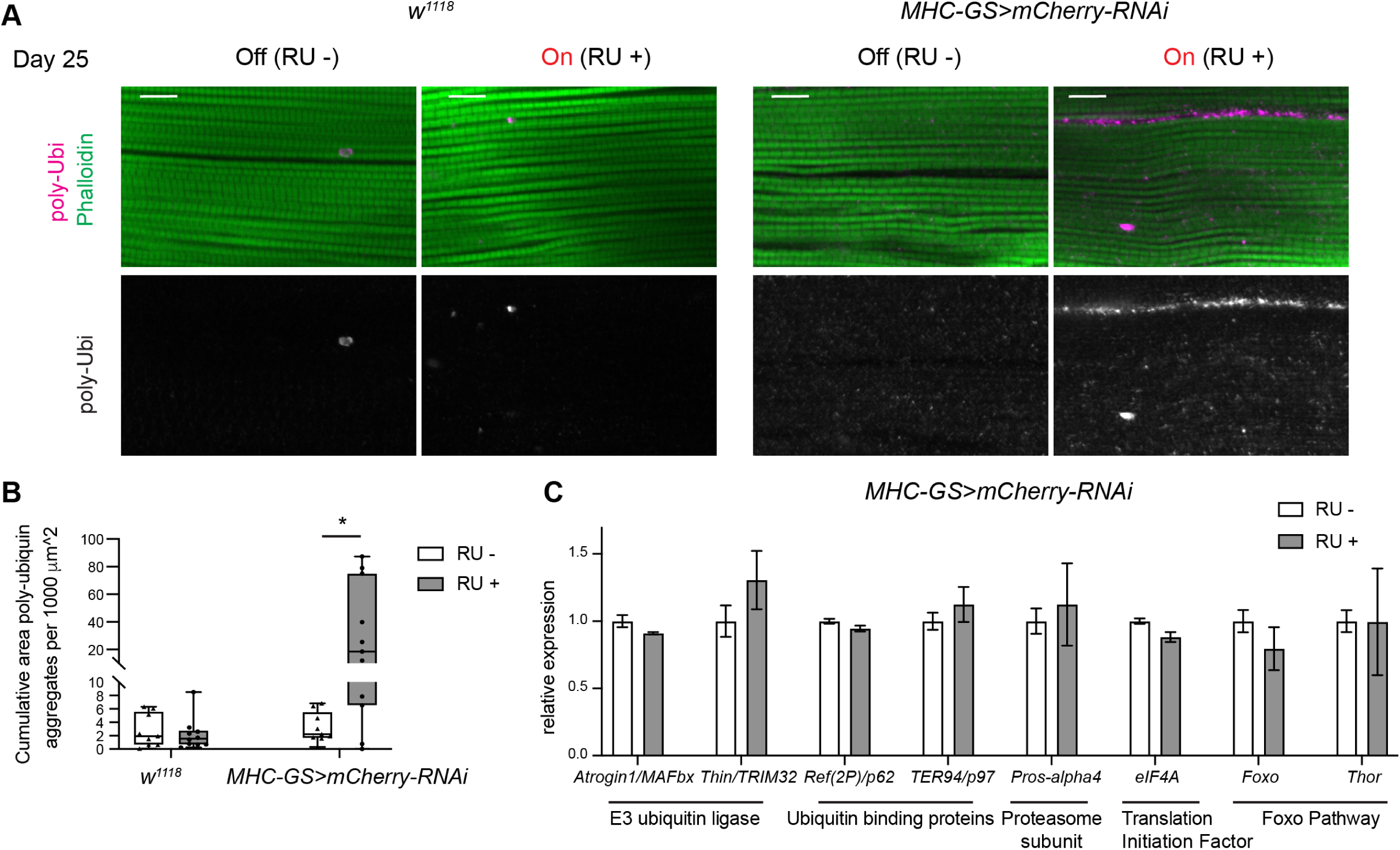
Formation of poly-ubiquitin aggregates in muscles is increased upon RU486-dependent activation of *MHC-GS-GAL4*. (A) Confocal section images of flight muscles in a sagittal view. Hemithorax sections of 25-d-old males stained with Phalloidin (green) and anti-poly-Ubi (magenta). Scale 10 µm. (B) Quantification of cumulative area of poly-ubiquitin aggregates. Box and whiskers, N = 9-10 hemithoraces/group for w1118 and N = 20-21 hemithoraces/group for *MHC-GS>mCherry-RNAi*. Mann Whitney test, * p<0.05. Note high disparity across our dataset, meaning that some animals showed very high levels of poly-ubiquitinated protein, while other show undetectable levels. (C) Gene expression measured by RT-qPCR in dissected thoraces of 25-day-old adult males. Fold change relative to control (RU - /Off group). Bar plots with bars, mean ± sem, N = 3 samples/group. Two-way ANOVA, p > 0.05, Bonferroni’s multiple comparisons test. Genotype: (A, B, C) *w*/Y*; +; *MHC-GS-GAL4*/*UAS-mCherry-RNAi*,(B) *w^1118^/Y*

In mammals, the ubiquitin-proteasome system is involved in the ubiquitination and protein degradation of skeletal muscles (Sandri *et al*. 2004; Cohen *et al*. 2012; Piccirillo and Goldberg 2012), yet, its role in *Drosophila* is not fully elucidated (Piccirillo *et al*. 2013). Therefore, we analyzed whether increased ubiquitination was the result of increased expression of genes encoding for components of the ubiquitin-proteasome system. RNA was extracted from thoraces of 25-day-old animals treated with RU+ and RU- and the expression of E3 ubiquitin ligases, such as *Atrogin1/MAFbx* and *Thin/TRIM32*, and the ubiquitin-binding proteins *Ref(2P)/p62* and *TER97/p97*, and the *proteasome 28kD subunit Pros-alpha4* was measured by RT-qPCR. As shown in Figure 3C, there were no significant differences in the mRNA levels of these genes between RU+/On and RU-/Off groups.

It was shown that activating FOXO and its downstream target Thor/4E-BP protects muscles from increased accumulation of polyubiquitin protein aggregates and proteostasis during aging (Demontis and Perrimon 2010). The mRNA expression of *FOXO* and *Thor/4E-BP* were not altered in animals treated with RU drug (Figure 3C), thus suggesting that the ubiquitination phenotype upon activation of the GeneSwitch system with RU486 is neither due to the FOXO pathway activation nor changes in the expression of the ubiquitin-proteasome system.

### Effect of exercising in RU486-dependent induction of *MHC-GS-GAL4* muscles

Regular exercise improves metabolism and numerous adaptive responses that contribute to a positive health benefit (Egan and Zierath 2013). In previous studies, the *MHC-GS-GAL4* driver has been used in chronic exercise model and the authors detected mostly an overall improvement in all assays when the GeneSwitch system was On (Kim *et al*. 2020). The physical endurance of flies is measured as a chronic exercise model and flies exhibit a robust response to exercise accompanied with a series of adaptations that improve overall physiology across various genotypes (Sujkowski and Wessells 2018; Damschroder *et al*. 2020). Given that exercising has a profound effect in muscle physiology we were wondering whether exercising can protect muscle tissue from the disruption seen in activated *MHC-GS-GAL4* flies. The 3-day-old *MHC-GS>mCherry-RNAi* flies were treated with RU + and RU -. Then, animals underwent a gradual training program that increases the training time once per week for three consecutive weeks (Figure 4A). We found that endurance at day 5, measured as time-to-fatigue and plotted as a survival curve (runspan), was already significantly reduced in RU + adult males (On, red lines) compared to their RU-control (Off, black lines, Figure 4B). Then, upon completing the training program, the exercised-trained flies (EX) experienced a series of adaptations, including increased endurance and increased climbing speed compared to age-matched controls for untreated flies (Off, Figure 4 C-D, EX, solid black line vs UN, dashed black line) (Tinkerhess *et al*. 2012; Sujkowski and Wessells 2018). However, turning On the GS system with *MHC-GS-GAL4* reduced the climbing endurance after three weeks of exercise training (runspan), as well as the acute climbing speed (Figure 4C-D, On EX, solid red line vs Off EX, solid black line). Intriguingly, the runspan in exercised-trained *MHC-GS>mCherry-RNAi* flies (On, EX, red solid line) was slightly longer than in their untrained age-matched flies (On, UN, red dashed line), which is an indication that exercise has a positive impact, even in this context. Furthermore, we confirmed that a similar response to exercise was detected using a second RNAi transgene, *MHC-GS>GFP-RNAi* (Figure 4E-G), thus further supporting the adverse effects caused by the activation of the *MHC-GS-GAL4* driver with RU486.

**Figure 4:**
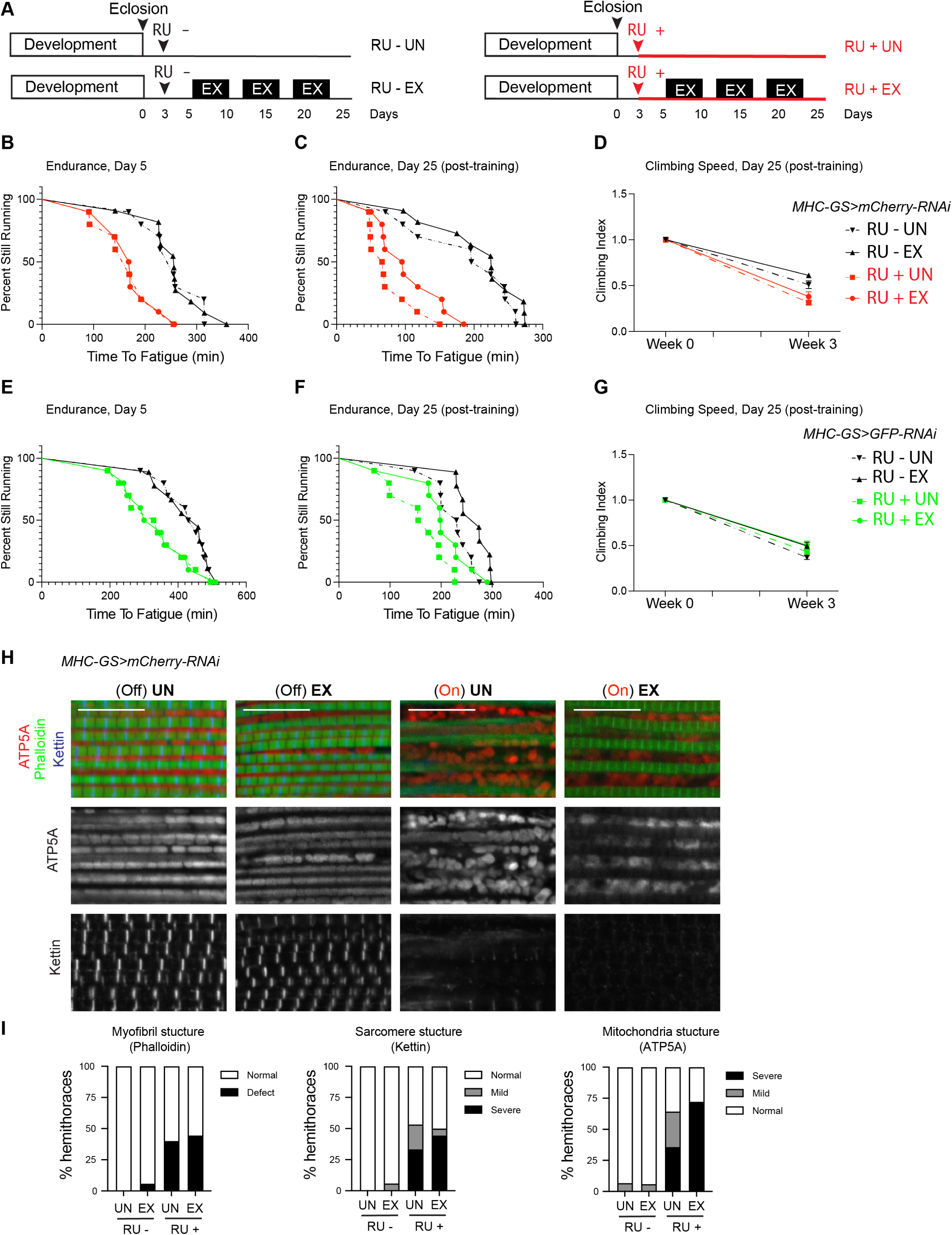
Chronic exercise does not protect muscles from defects induced by RU486-dependent activation of *MHC-GS-GAL4*. (A) Schematic showing time of drug treatment and times of exercise training. Exercise took place shortly after lights-on and lasted for 1.5 hours in week 1, 2 hours in week 2, and 2.5 hours in week 3 for 5 days each week. (B) Young 5-day old flies expressing *mCherry-RNAi* with the *MHC-GS-GAL4* driver ran significantly shorter than genetically identical flies without the inducing RU486 drug (log-rank, p<.001) prior to receiving any exercise treatment. (C) Age-matched 25-day-old flies expressing *mCherry-RNAi* with the *MHC-GS-GAL4* driver ran significantly shorter than genetically identical flies without the inducing RU486 drug (log-rank, p<.001), regardless of exercise treatment. Exercise did improve endurance significantly in both RU+ and RU-groups (log-rank, p < 0.001). (D) Climbing speed was consistently lower in flies expressing *mCherry RNAi* with the *MHC-GS-GAL4* driver than in controls without the inducing RU486 drug, when measured longitudinally across 25 days of age (2-way ANOVA, p<.001). Exercise did not reverse this defect. (E) Young 5-day old flies expressing *GFP-RNAi* with the *MHC-GS-GAL4* driver ran significantly shorter than genetically identical flies without the inducing RU486 drug (log-rank, p<.001) prior to receiving any exercise treatment. (F) Age-matched 25-day-old flies expressing *GFP-RNAi* with the *MHC-GS-GAL4* driver ran significantly shorter than genetically identical flies without the inducing RU486 drug (log-rank, p<.001), regardless of exercise treatment. Exercise did improve endurance significantly in both RU+ and RU-groups (log-rank, p < 0.001). (G) Climbing speed was not significantly altered across 25 days of longitudinal measurement by expression of *GFP-RNAi* with the *MHC-GS-GAL4* driver, (2-way ANOVA, p > 0.05). (H) Confocal section images of flight muscles in a sagittal view. Hemithorax sections of 25-d-old males stained with Phalloidin (green), anti-kettin (blue), and anti-ATP5A (red). Scale 10 µm. (I) Quantification of percentage of hemithoraces displaying either normal or defective (mild and sevre) morphology for (left) overall myofribril structure, (middle) sarcomere structure, and (right) mitochondria shape (i.e., round). Stacked bars, N = 14-18 hemithoraces/group. Genotypes: (B-D, H,I) *w*/Y*; +; *MHC-GS-GAL4*/*UAS-mCherry-RNAi*, (E-G) *w*/Y*; *UAS-GFP-RNAi*/+; *MHC-GS-GAL4*/+.

Both exercised-trained and untrained *MHC-GS>mCherry-RNAi* animals were harvested at 25-days-old to analyze overall structure of their flight muscles. When GS system was Off, the overall muscle organization of exercised animals were indistinguishable from untrained animals (Figure 4H). However, in animals fed on RU drug diet, in which gene switch system was turned On, again we found disrupted muscles (Figure 4H-I). Overall, our data imply that despite the beneficial effect of exercising, chronic exercise does not prevent muscle disruption in adult flies when inducing the RU486-dependent *MHC-GS-GAL4* driver.

## Discussion

The RU486-dependent GeneSwitch system is a valuable tool because it expands the application of the conventional GAL4/UAS system by adding a temporal regulation to the expression of the tissue specific transgene. Therefore, this inducible system has been widely used with numerous GS-GAL4 drivers. Here, we report that turning on *MHC-GS-GAL4* driver alone without a UAS transgene results in alteration of the muscle structure, and in severe reduction of animal lifespan. Our findings show that this is not a result of a response to RU486 since feeding the wild type strain *w^1118^* with RU486 supplemented food produced no phenotype. Rather we suggest that these unwanted effects are due to turning On the *MHC-GS-GAL4* driver with RU486. Thus, caution should be exercised when using this system as it can lead to erroneous data interpretation. One conclusion from the data reported here is that the comparison between induced and non-induced groups using the GeneSwitch system are insufficient. As the effects may vary depending on the stock and tissue under investigation, we recommend setting up additional control groups to validate conclusions for every experimental condition, which is in line with other reports (Poirier *et al*. 2008; Scialo *et al*. 2016). Additionally, a mock UAS-RNAi transgene, such as *mCherry-RNAi* or *eGFP-RNAi*, should be included to account for any adverse phenotype induced by the GeneSwitch system on its own.

In a recent study, defects in muscle function and mitochondrial gene expression have also been reported when inducing the GeneSwitch system in adult muscles with both *MHC-GS-GAL4* and *Act88F-GS*-*GAL4* drivers (Robles-Murguia *et al*. 2019), which is in agreement with our study. Interestingly, defects were mostly associated with these muscle-specific drivers but not other wild type strain nor other GS-GAL4 drivers, thus confirming that the effects vary depending on the stock and tissue explored. We noticed that the earliest phenotype that we found was defects in mitochondrial structure in 5-day old flies suggesting that non-functional mitochondria may contribute to the disruption of myofibrils and sarcomere units. This is further supported by the fact that myofibrils and mitochondria morphogenesis are intimately linked during development (Avellaneda *et al*. 2021).

Noticeably, the *MHC-GS-GAL4* has been successfully used in other studies exploring different phenotypes and transgenes (Kim *et al*. 2020). Sestrin was overexpressed in muscles using *MHC-GS*-*GAL4* and it led to both molecular and physiological effects of exercise, and consequently extended endurance (Kim *et al*. 2020). Since endurance exercise training provides numerous health benefits, including increasing mitochondrial biogenesis and activity (Holloszy and Booth 1976), we reasoned that it could block muscle disruption induced by *MHC-GS-GAL4*. However, we found no significant response to endurance exercise training in RU486-induced expression of mock-RNAi transgenes. Altogether, these data suggest that the effects of activated *MHC-GS-GAL4* are context-dependent and may vary based on tissue, construct being expressed, and/or other factors. In another study, no defective lifespans were found in mock-treated control flies when comparing between RU486-treated and untreated groups (*MHC-GS*>+, RU-vs RU+) (Hunt and Demontis 2022), which is in contrast with what we found. One potential explanation for the difference is that RU486 was used at 50μM concentration in this study, which is four times less than what we and others have used (McGuire *et al*. 2004). Thus, variation in the extent of the adverse effects may also depend on the exact concentration of RU486 to activate the GeneSwitch system with the *MHC-GS-GAL4* driver. As most reported negative effects from RU-activated *MHC-GS-GAL4* have been associated with expression of RNAi transgenes, it is possible that RNAi transgenes warrant particular care with this driver system.

The main conclusion of our work is that inducing *MHC-GS-GAL4* with RU486 leads to adverse effects in mitochondrial organization, adult skeletal muscle structure, and ultimately results in shorter lifespan. Hence, caution should be taken when using this inducible system to avoid erroneous data interpretation. The adverse effect of activating RU486-dependent *MHC-GS-GAL4 per se* varies upon the concentration of RU486 (50μM vs 200μM), the type of transgene (knockdown *vs* overexpression), and the developmental stage (larval muscles *vs* adult muscles). Thus, we recommend setting up an additional control by analyzing the effect of RU486-induced expression of a mock transgene when using the GeneSwitch system instead of relying on only comparing the phenotypes between RU486-treated and untreated animals.

## Materials and Methods

### Fly maintenance and stocks

Fly stocks were maintained on standard cornmeal food at 25°C. The flies *w^1118^* (BDSC 3605), *UAS-mCherry-RNAi* (P{VALIUM20-mCherry}attP2, BDSC 35785), *UAS-GFP-RNAi* (P{UAS-GFP.dsRNA.R}143, BDSC 9331) and *MHC-GS-GAL4* (P{Mhc-Switch.O}GSG314-2, BDSC 43641) were used.

### Activation of GeneSwitch system

The compound RU-486 (Mifepristone, Sigma, M8046) was prepared as 10mM stock in 70% EtOH. Standard cornmeal food was melted in microwave and RU486 was added to a final concentration of 0.2 mM (RU+). In parallel 70% EtOH was added to vehicle vials labeled as RU-. Crosses were set up in normal conditions and maintained at 25°C. On day 2 after eclosion between 15 and 25 males were collected and transferred to vials with standard cornmeal food supplemented with either 0.2 mM RU386 (RU+) to turn On the system or vehicle (70% ethanol, RU-, system Off). Flies were maintained at 25°C and transferred to fresh RU+ and RU-vials twice a week until harvested.

### RT-qPCR

Thoraces with legs from a total of 10 males were collected per sample and homogenized using 1 mL TRIzol (Invitrogen #15596026). Total RNA was quantified using Qubit RNA HS assay kit. About 400ng total RNA was used in SensiFast cDNA synthesis kit (Bioline, BIO-65053). The RT-qPCR reactions were set up using SensiFAST SYBR No-ROX kit (Bioline, BIO-98005) in a Light Cycler 480 II system (Roche). Data is shown as relative expression normalized to geometric average of RpL30 and RpL32. Two technical replicates and three biological replicates were included. Primer sequences are listed in Supplemental Table S1.

### Immunostaining and confocal imaging

About ten males per group were collected. Thoraces were dissected and fixed in 4% formaldehyde in relaxing buffer (20 mM Sodium phosphate, 5 mM MgCl2, 0.1% Triton-X, 5mM EGTA, 5 mM ATP) for 20 min (Weitkunat and Schnorrer 2014). Then, tissue was sectioned in the sagittal plane, and incubated in the fixation buffer for an additional 15 min. Hemi-thoraces were incubated in 0.3% Triton X-100 in PBS, then in 2% bovine serum albumin (BSA) 0.3% Triton X-100 in PBS for 1 hour and with primary antibodies overnight at 4°C. A total of four washes were done in 0.3% Triton X-100 followed by incubating with secondary antibody for 2 hours in 10% normal goat serum and 0.1% Triton X-100 in PBS. Tissue was washed four times, and stored in glycerol with antifade at 4°C. All steps were carried out at room temperature with gentle agitation otherwise indicated.

The primary antibodies were anti-ATP5A (abcam, ab14748, mouse, 1:500), anti-Kettin (MAC155, rat, 1:1000, Babraham Institute) and anti-Ubiquitin mAb (clone FK2, MilliporeSigma, ST1200). The secondary antibodies used are Alexa Fluor 647 anti-Rat IgG and Cy3 anti-Mouse IgG (Jackson Immunoresearch, 712-605-153 and 715-165-151). The dyes Phalloidin-Atto-488 (Sigma-Aldrich, 49409, 1:500,) and DAPI (1:500) were used.

Laser Scanning Microscope 700 (Zeiss LSM700, Observer.Z1) was used to acquire fluorescent images with objectives x20/0.8, x40/1.2 and x100/1.45. Among the indirect flight muscles the dorsal longitudinal muscles were examined here. All images are confocal single-plane images and representative images were processed using Photoshop (Adobe Systems).

### Endurance

Vials of 20 flies each were placed on an automated climbing machine known as the Power Tower (Piazza *et al*. 2009) and exposed to continuous negative geotaxis stimulus, until they became too fatigued to respond ((Damschroder *et al*. 2018). Fatigue was defined as a lack of response from 80% of flies in a vial for three consecutive stimulus inductions. Vials were analyzed as single statistical units and time-to-fatigue was plotted and analyzed by log-rank. Significance was set at p<0.05.

### Climbing speed

Speed was measured as previously described (Damschroder *et al*. 2018). Briefly, vials of 20 flies were manually tapped down to induce negative geotaxis and photographed to measure progress up the vial after 2 seconds. Position of flies in the vial was analyzed by Image J and averaged across 4 repetitions for each vial per timepoint. Measurements were taken longitudinally across time using the same cohort. Data was analyzed using 2-way ANOVA with a post-hoc Bonferroni correction for multiple measurements.

### Survival curve

Between 15 and 25 males per vial were maintained at 25°C in either RU+ or RU-food. Flies were directly flipped to fresh vials twice a week. Dead flies were scored daily around the time *GMH-GS-GAL4* flies were dying between 10 and 17 day-old. Data was plotted as Kaplan-Meier plot, a total of three vials per group were used.

### Quantification and statistical analysis

All images were blinded and randomized prior to quantification. One image per hemithorax was taken and analyzed. The overall structure of the myofibrils was scored as either normal or defective based on stained with phalloidin. The overall structure of the sarcomere was scored as either normal, mild or severe phenotype based on the staining with anti-kettin. The overall shape of mitochondria was scored as normal, mild or severe phenotype based on the staining with anti-ATP5A. The percentage for each group in displayed on the graphs.

The total area of poly-ubiquitin aggregates was quantified using the ‘analyze particles’ function of ImageJ. For each image the region of interest was selected randomly based on phalloidin channel. Cumulative area was normalized to the total area of region of interest.

Statistical analysis was performed using GraphPad Prism v10 (GraphPad Software). Details on the sample size, number of independent experiment, and statistical analysis are listed in figure legends.

## Data availability

All data are included in this article and its supplementary information files.

## Acknowledgments

We thank Carthic Rajagopalan and Isabel Liseth for technical support. We thank the Bloomington Drosophila Stock Center (NIH P40OD018537) for providing the stocks. We are grateful to Flybase for online resources on the Database of Drosophila Genes and Genomes.

## Author contribution

Conceptualization, M.P.Z., D.D, R.J.W. and M.V.F.; Investigation, M.P.Z, D.D, A.W. and C.R.; Supervision, M.P.Z., R.J.W., and M.V.F.; Visualization, M.P.Z.; Writing – Original Draft, M.P.Z. and M.V.F.; Writing – Review & Editing, M.P.Z., R.J.W. and M.V.F; Funding Acquisition, M.V.F., R.J.W.

## References

Avellaneda, J., C. Rodier, F. Daian, N. Brouilly, T. Rival et al., 2021 Myofibril and mitochondria morphogenesis are coordinated by a mechanical feedback mechanism in muscle. Nat Commun 12: 2091.

Brand, A. H., and N. Perrimon, 1993 Targeted gene expression as a means of altering cell fates and generating dominant phenotypes. Development 118: 401–15.

Damschroder, D., T. Cobb, A. Sujkowski, and R. Wessells, 2018 Drosophila Endurance Training and Assessment of Its Effects on Systemic Adaptations. Bio Protoc 8: 1–19.

Damschroder, D., K. Richardson, T. Cobb, and R. Wessells, 2020 The effects of genetic background on exercise performance in Drosophila. Fly (Austin) 14: 80–92.

Demontis, F., and N. Perrimon, 2010 FOXO/4E-BP signaling in Drosophila muscles regulates organism-wide proteostasis during aging. Cell 143: 813–25.

Egan, B., and J. R. Zierath, 2013 Exercise metabolism and the molecular regulation of skeletal muscle adaptation. Cell Metab 17: 162–184.

Haas, K. F., E. Woodruff, and K. Broadie, 2007 Proteasome function is required to maintain muscle cellular architecture. Biol Cell 99: 615–626.

Holloszy, J. O., and F. W. Booth, 1976 BIOCHEMICAL ADAPTATIONS TO ENDURANCE EXERCISE IN MUSCLE. Annu. Rev. Physiol. 38: 273–291.

Hunt, L. C., and F. Demontis, 2022 Age-Related Increase in Lactate Dehydrogenase Activity in Skeletal Muscle Reduces Life Span in Drosophila. Journals of Gerontology - Series A Biological Sciences and Medical Sciences 77: 259–267.

Kim, M., A. Sujkowski, S. Namkoong, B. Gu, T. Cobb et al., 2020 Sestrins are evolutionarily conserved mediators of exercise benefits. Nat Commun 11: 1–14.

McGuire, S. E., Z. Mao, and R. L. Davis, 2004 Spatiotemporal gene expression targeting with the TARGET and gene-switch systems in Drosophila. Science’s STKE pl6.

Osterwalder, T., K. S. Yoon, B. H. White, and H. Keshishian, 2001 A conditional tissue-specific transgene expression system using inducible GAL4. Proc Natl Acad Sci U S A 98: 12596– 12601.

Piazza, N., B. Gosangi, S. Devilla, R. Arking, and R. Wessells, 2009 Exercise-training in young Drosophila melanogaster reduces age-related decline in mobility and cardiac performance. PLoS One 4: e5886.

Poirier, L., A. Shane, J. Zheng, and L. Seroude, 2008 Characterization of the Drosophila Gene-Switch system in aging studies: A cautionary tale. Aging Cell 7: 758–770.

Robles-Murguia, M., L. C. Hunt, D. Finkelstein, Y. Fan, and F. Demontis, 2019 Tissue-specific alteration of gene expression and function by RU486 and the GeneSwitch system. NPJ Aging Mech Dis 5: 2–6.

Roman, G., K. Endo, L. Zong, and R. L. Davis, 2001 P{switch}, a system for spatial and temporal control of gene expression in drosophila melanogaster. Proc Natl Acad Sci U S A 98: 12602–12607.

Scialo, F., A. Sriram, R. Stefanatos, and A. Sanz, 2016 Practical recommendations for the use of the geneswitch Gal4 system to knock-down genes in drosophila melanogaster. PLoS One 11: 1–13.

Sujkowski, A., and R. Wessells, 2018 Using Drosophila to Understand Biochemical and Behavioral Responses to Exercise. Exerc Sport Sci Rev 46: 112–120.

Tinkerhess, M. J., S. Ginzberg, N. Piazza, and R. J. Wessells, 2012 Endurance training protocol and longitudinal performance assays for Drosophila melanogaster. J Vis Exp 1–6.

Weitkunat, M., and F. Schnorrer, 2014 A guide to study Drosophila muscle biology. Methods 68: 2–14.

